# Securing diagonal integration of multimodal single-cell data against ambiguous mapping

**DOI:** 10.1101/2023.10.05.561049

**Authors:** Han Zhou, Kai Cao, Yang Young Lu

## Abstract

Recent advances in single-cell multimodal omics technologies enable the exploration of cellular systems at unprecedented resolution, leading to the rapid generation of multimodal datasets that require sophisticated integration methods. Diagonal integration has emerged as a flexible solution for integrating heterogeneous single-cell data without relying on shared cells or features. However, the absence of anchoring elements introduces the risk of artificial integrations, where cells across modalities are incorrectly aligned due to ambiguous mapping. To address this challenge, we propose SONATA, a novel diagnostic method designed to detect potential artificial integrations resulting from ambiguous mappings in diagonal data integration. SONATA identifies ambiguous alignments by quantifying cell-cell ambiguity within the data manifold, ensuring that biologically meaningful integrations are distinguished from spurious ones. It is worth noting that SONATA is not designed to replace any existing pipelines for diagonal data integration; instead, SONATA works simply as an add-on to an existing pipeline for achieving more reliable integration. Through comprehensive evaluation on both simulated and real multimodal single-cell datasets, we observe that artificial integrations in diagonal data integration are widespread yet surprisingly overlooked, occurring across all mainstream diagonal integration methods. We demonstrate SONATA’s ability to safeguard against misleading integrations and provide actionable insights into potential integration failures across mainstream methods. Our approach offers a robust framework for ensuring the reliability and interpretability of multimodal single-cell data integration. ^1^

## 1 Introduction

Recent advances in single-cell multimodal omics technologies now allow biologists to simultaneously capture unprecedented high-resolution omics profiles at the cellular level [1]. Numerous technologies have been developed to explore cellular systems from diverse perspectives, resulting in multiple data modalities, each capturing a distinct molecular aspect of the cell. These modalities include scRNA-seq for gene expression [2], scATAC-seq for chromatin accessibility [3], scMethyl-seq for DNA methylation [4], scHi-C for chromatin 3D conformation [5], and scChIC-seq for histone modifications [6], and others. Consequently, the integrative analysis of this vast single-cell multimodal omics data has garnered increasing interest, with the potential to provide deeper insights into cellular processes that were previously inaccessible through studies of bulk cells.

Despite its great potential, integrating single-cell data modalities for downstream interpretation remains a complex challenge [7]. First of all, different data modalities may not share the same set of features. For example, scRNA-seq focuses on gene expression, while scATAC-seq measures chromatin accessibility across various genomic regions. Beyond feature discrepancies, another challenge arises from the destructive nature of profiling technologies, which results in different data modalities measuring distinct sets of cells. For example, both scATAC-seq and scHi-C target genomic DNA, meaning that each individual cell can only be measured once due to genome cleavage. Although recent technological advances allow for joint profiling of some modalities in the same cells (such as scRNA-seq and scATAC-seq), many modalities still cannot be measured together. Additionally, many single-cell datasets have already been collected over the years using distinct sets of cells. Therefore, maximizing the utility of these datasets through integration is essential, rather than disregarding valuable data collected with distinct cell sets. Lastly, intrinsic variations exist between datasets, even within the same modality. These variations can be biological, arising from samples collected under different conditions across tissues, organs, individuals, or species. They can also be technical, resulting from differences in laboratory instruments and experimental protocols used during sample profiling. In summary, the absence of shared cells or features across heterogeneous single-cell modalities makes data integration a highly challenging task.

In recent years, a growing number of integration methods have been developed to tackle multimodal single-cell data integration, using different approaches depending on the availability of shared features or shared cells as anchors [8]. Among these methods, the “diagonal integration” approach—the focus of this paper—has garnered increasing attention, as it does not rely on anchoring cells or features for integration, making it the most flexible option with minimal prior knowledge. In contrast, methods that rely on anchored features or cells are only applicable when it is possible to engineer matched features or when multiple modalities can be jointly profiled within the same cell. From a practitioner’s perspective, an effective diagonal integration method would significantly broaden the scope of potential data integration, making it highly appealing to the community. However, the absence of anchors also introduces greater challenges in achieving accurate integration results [9]. Grounded in empirical evidence suggesting that each data modality occupies a low-dimensional manifold with latent semantic structure [10], we note that diagonal integration methods, including MMD-MA [11], SCIM [12], SCOT [13, 14], UnionCom [15], and Pamona [16], typically involve three steps (Fig. 1**a**): (1) Projecting different modalities into a shared space to reveal their underlying manifold structures, (2) Identifying correspondence relationships between cells across data modalities, and (3) Generating an integrated profile from all modalities. Although the methods differ in implementation details, they all share the same principle of aligning data within a low-dimensional manifold [9].

**Figure 1.**
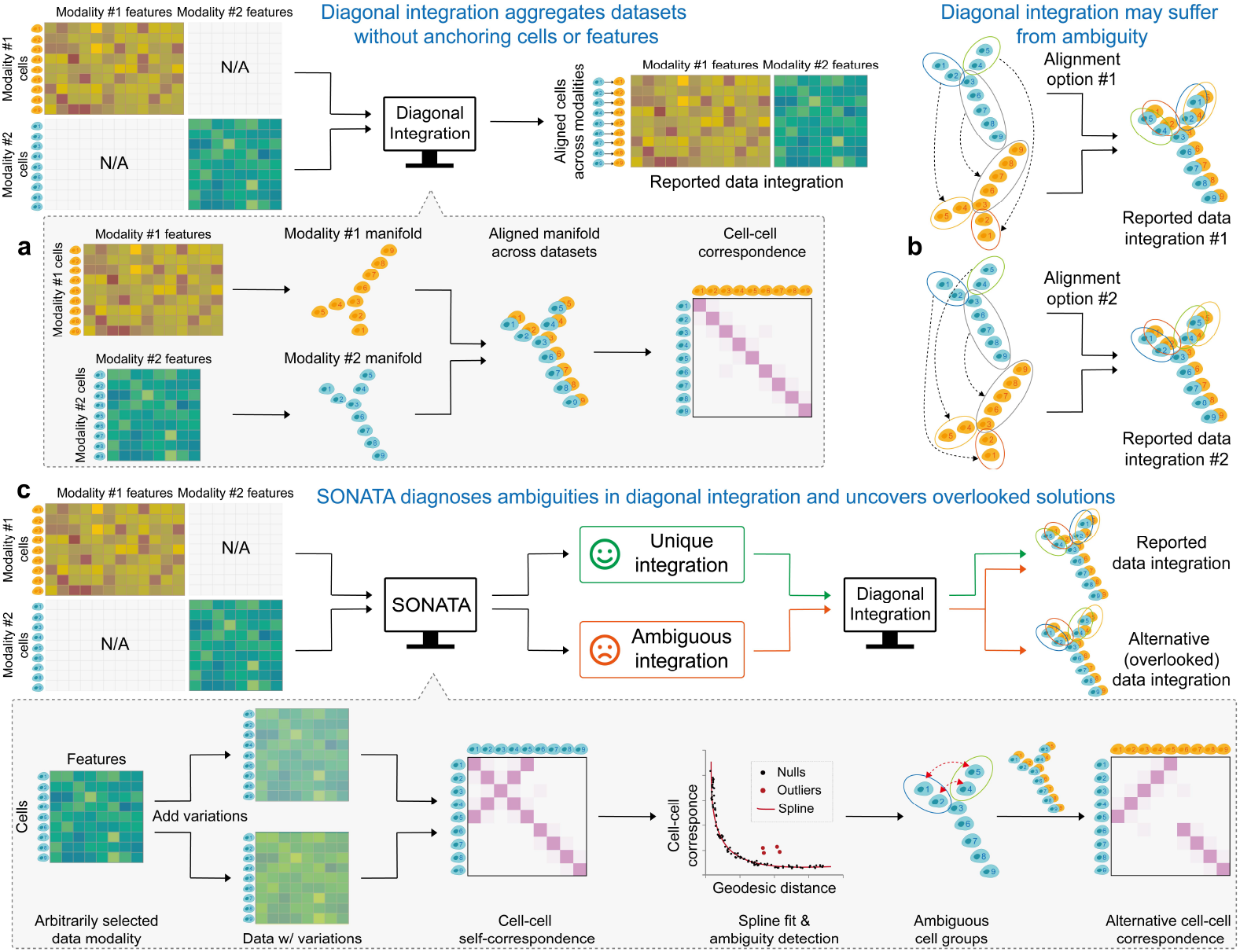
Overview of SONATA. (**a**) SONATA is designed to safeguard diagonal integration of multimodal single-cell data, an approach that does not depend on anchoring cells or features for integration. (**b**) SONATA identifies potential artificial integrations arising from ambiguous mapping, which occurs when two cells that are far apart in the data manifold share similar geometric contexts. (**c**) SONATA uses a novel cell-cell ambiguity measurement to provide insights into possible alternative integration solutions that are often overlooked by existing integration methods. The ambiguity is measured using a statistical confidence measure for each pair of cells within the same modality. This ambiguity is then used to identify cell groups with similar geometric contexts, which could potentially lead to artificial integrations in cross-modality mapping.

Despite the varied implementations of manifold alignment, all existing diagonal integration methods share an implicit yet critical assumption: data from different modalities are generated from a similar distribution, as maintained by the shared manifold [9]. As a consequence, these methods indiscriminately provide an integration solution along with corresponding cross-modality correspondence relationships. As a user, a natural question arises: “how can one distinguish between true biological integration—accurately matching cells across different modalities—and any potential artificial integration?” Specifically, an artificial integration solution can be mathematically indistinguishable from the accurate biological solution we seek. Studies have reported that multiple alignments can appear equally optimal, potentially leading to artificial alignments [17]. As a motivating example, consider the cell cycle [18], a periodic biological process in human cells. The periodicity of cycling cells in different transcriptomic states forms a circular trajectory, with a cell’s position on the trajectory indicating its timing in the cycle. In this case, an infinite number of incorrect mappings can exist between two cycles, as rotating one cycle by arbitrary degrees naturally produces new, albeit incorrect, solutions. Without a systematic mechanism to flag potential artificial integrations, blindly integrating single-cell datasets can lead to unfaithful and misleading interpretations [19].

In this paper, we propose a novel diagnostic method for diagonal data integration of multimodal single-cell data, SONATA (Securing diagOnal iNtegrATion against Ambiguous mapping), that aims to identify potential artificial integrations resulting from ambiguous mapping (Fig. 1). Here, ambiguous mapping does not refer to substituting one cell with another within the same modality due to their similarity in features or close proximity in the underlying manifold. Instead, it occurs when two cells that are far apart in the data manifold share similar geometric contexts, leading to potential confusion during integration. For example, as illustrated in Fig. 1**b**, cell 1 in data modality 1 is expected to map to cell 1 in data modality 2. Ambiguous mapping refers to the scenario where cell 1 in data modality 1 is instead mapped to cell 5 in data modality 2. It’s important to note that mapping cell 1 in data modality 1 to cell 2 in data modality 2 is not considered ambiguous; this could simply result from variations in the data or the inherent instability of diagonal integration methods. Unlike existing methods that recklessly report an arbitrary integration solution, SONATA distinguishes whether the integration is unique or if multiple alternative solutions exist, prompting users to carefully discern the true biological solution before proceeding to downstream analysis.

The novelty of SONATA are twofold. First and foremost, we demonstrate that artificial integrations resulting from ambiguous mapping in diagonal data integration are widespread yet surprisingly overlooked, occurring across all mainstream diagonal integration methods. Note that artificial integrations are more harmful than failed integrations because, while failed integrations can be qualitatively recognized, artificial integrations are difficult to detect and can mislead users into pursuing hypotheses based on erroneous results. For this reason, we propose a systematic strategy to detect potential artificial integrations without relying on prior knowledge (Fig. 1**c**). The core idea is centered on a novel cell-cell ambiguity measurement, where two cells are considered ambiguous—and thus likely to substitute for each other in a cross-modality alignment—given that they are far apart in the data manifold. By aligning a data modality to a variational version of itself, ambiguity is assessed through a statistical confidence measure for each pair of cells within the same modality. This is based on their observed correspondences relative to a null model, in which the correspondence probability diminishes as the cell-cell distance increases along the data manifold. Consequently, the cell-cell ambiguities provide insights into possible alternative integration solutions that are often overlooked by existing integration methods.

We applied SONATA to both simulated and real multimodal single-cell datasets to demonstrate its empirical utility. Our experiments on real datasets highlight SONATA’s ability to safeguard diagonal integration of gene expression, DNA methylation, and chromatin accessibility modalities against ambiguous mapping from mainstream methods, effectively informing potential users where these methods might fail. It is worth noting that SONATA is not designed to replace any existing pipelines for diagonal data integration; instead, SONATA works simply as an add-on to an existing pipeline for achieving more reliable integration. In summary, SONATA’s diagnostic analysis is a pioneering effort that offers guidelines for the reliable use of diagonal integration methods, ensuring robust and interpretable integration of single-cell data.

### 1.1 Related work

Most diagonal integration methods rely on unsupervised manifold alignment [9], a technique designed to discern a low-dimensional manifold that captures covariation across various data modalities in the absence of cross-modality correspondence information. As one of the pioneering efforts, MATCHER [10] employs a Gaussian process latent variable model for integration, albeit constrained to aligning 1-D trajectories. Recent methods, such as MMD-MA [11], SCIM [12], SCOT [13, 14], UnionCom [15], and Pamona [16], have extended the capabilities of MATCHER to accommodate more intricate structures. When cross-modality correspondence information is readily available, diagonal integration simplifies to horizontal or vertical integration [7], depending on the presence of shared features or cells, respectively. In horizontal integration, features between modalities exhibit one-to-one correspondence and serve as anchors for integrating the datasets; batch correction methods fall into this category, including Harmony [20], Scanorama [21], Seurat [22], LIGER [23], and so on. For example, Seurat [22] employs canonical correlation analysis to identify shared manifold space, while LIGER [23] achieves integration through integrative non-negative matrix factorization. In vertical integration, multiple modalities are concurrently profiled from the same samples, and the datasets are integrated by concatenating the features, including Hetero-RP [24], MOFA+ [25], iNMF [26], and so on. For example, Hetero-RP [24] concatenates and rescales features from different modalities, prioritizing important features with higher weights than others.

## 2 Approach

### 2.1 Problem setup

Let the two datasets of cells be 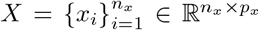 in data modality X and 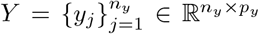 in data modality 𝒴, respectively. The numbers of cells in the two data modalities are *n*_*x*_ and *n*_*y*_, and the feature dimensions are *p*_*x*_ and *p*_*y*_, respectively. Given *X* and *Y* as input, existing diagonal integration methods uncover a low-dimensional manifold that captures the covariation between *X* and *Y* and identify correspondence relationships between cells across the modalities (Fig. 1**a**). Each data modality is typically represented as a low-dimensional manifold by constructing a geodesic distance matrix between the cells. To achieve this, a weighted *k*-nearest neighbor (*k*-NN) graph of cells within each data modality is constructed [27]. Subsequently, the shortest distance between each node pair on the graph is calculated, as these shortest distances serve as approximations of geodesic distances on the data manifold [28]. The resultant geodesic matrices for *X* and *Y* are denoted as 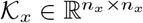 and 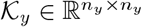, respectively.

Once two datasets are aligned, the identified correspondence relationships are encoded into a crossmodality correspondence matrix 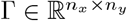, which is generated either explicitly [13, 15] or implicitly [11, 12], where Γ_*ij*_ represents the probability of the *i*-th cell in χaligning with the *j*-th cell in 𝒴. SONATA focuses on diagnosing diagonal data integration by assessing whether the resulting integration may be artificial. If so, SONATA is designed to enumerate all possible solutions that may have been overlooked, ensuring that the true biological solution is included among the possibilities.

### 2.2 Measuring the manifold ambiguity between cells

SONATA takes an arbitrarily selected data modality as input and identifies ambiguous cells which could potentially lead to artificial integrations in cross-modality mapping. We reason that if such ambiguity exists, it indicates that certain regions of the data manifold resemble others, implying that these regions should align when the data is mapped against itself. However, utilizing self-alignment of the given data modality for ambiguity detection is challenging, as a trivial solution always exists where each cell aligns with itself rather than with geometrically similar cells that are far apart in the data manifold. To prevent a trivial solution, we align the data modality to a variational version of itself by introducing random Gaussian noise and varying the number of neighbors in manifold construction, making the alignment process more challenging (Fig. 1**c**). (Refer to Sec. A.3 for the details to construct variational data).

With the variations of the data modality, we repurpose existing diagonal integration methods to perform self-alignment, ignoring the self-correspondence between cells. In principle, any existing method can be applied here; in this paper, we use SCOT [13] because it explicitly reports a probabilistic cell-cell correspondence, making it convenient for subsequent identification of ambiguous cells. The resultant cell-cell correspondences from self-alignment are utilized to assess manifold ambiguity between cells using a statistical confidence measure. Specifically, SONATA models an empirical null from the observed cell-cell correspondences, by observing how it diminishes conditioned on the geodesic distance between cells along the data manifold. To achieve this, SONATA fits a cubic smoothing spline to model the probability of cell-cell correspondence as a function of geodesic distance, with an anti-tonic regression constraint to ensure the probability is monotonically non-increasing as the distance increases (Fig. 1**c**). (Refer to Sec. A.2.1 for the details to facilitate smooth and efficient spline fit). Once the spline is fitted, we calculate the deviation between the observed and expected correspondence probabilities for each cell pair. A one-sided p-value is then computed for each deviation using a normality test to evaluate its statistical significance, with a cutoff threshold set at 1%. A statistically significant ambiguous cell pairs indicates that these two cells share similar geometric contexts, potentially leading to artificial integrations in cross-modality mapping.

### 2.3 Diagnosing diagonal integration solutions

The presence of ambiguous cell pairs suggests the existence of alternative yet overlooked integration solutions. The existence of multiple integration solutions suggests that the solution provided by current diagonal integration methods may be artificial. Based on the intuition that cells engage in ambiguous alignments alongside their neighboring cells rather than individually, we apply a data-driven approach to aggregate ambiguous cells into more coherent and interpretable groups. Specifically, the aggregation of ambiguous cell pairs is framed as a constrained clustering problem [29], extending standard clustering by incorporating pairwise ambiguity constraints, which ensure that two cells identified as ambiguous cannot be placed within the same larger group. The final number of ambiguous groups is selected using the Elbow method, where the number of violated constraints is plotted against the number of groups. As the number of groups increases, the constraint violations decrease, and the optimal group number is determined at the point where this decrease slows and begins to plateau. The rationale for these ambiguous cell groups is that one group can be substituted for another based on their geometric resemblance during cross-modality integration.

Lastly, SONATA generates alternative integration solutions based on identified ambiguous cell groups (Fig. 1**c**). (Refer to Sec. A.2.2 for the details of alternative solution generation). It is important to note that the alternative integration solutions identified are based on geometric resemblance rather than biological validity. However, revealing these multiple possibilities encourages users to critically evaluate and discern the true biological solution, rather than relying on a potentially misleading outcome.

## 3 Materials and methods

### 3.1 Datasets

To uncover the artificial integration issues caused by ambiguous mapping, we first constructed four simulated datasets (Fig. 2**a**). The simulated datasets fall into two categories: one without ambiguity and three with ambiguity. The ambiguous datasets consist of T-shaped, Y-shaped, and X-shaped branches, designed to mimic bifurcating or multifurcating differentiation processes. The distinction among these datasets lies in the resemblance of two, three, and four branches, respectively. In contrast, the unambiguous dataset features a hook-like shape with a decaying density from one end to the other, where two manifolds are uniquely aligned without any ambiguity. Each simulated dataset comprises two modalities, each containing *n*_*x*_ = *n*_*y*_ = 300 cells, with cell-cell correspondence information generated. The cells reside on a 2-dimensional manifold for both modalities, which are then mapped to feature spaces of *p*_*x*_ = 1000 and *p*_*y*_ = 2000 dimensions, respectively, with Gaussian noise (*σ* = 0.1) added to the features. Given the differing feature dimensions between the two modalities, no feature-level correspondence exists between them.

**Figure 2.**
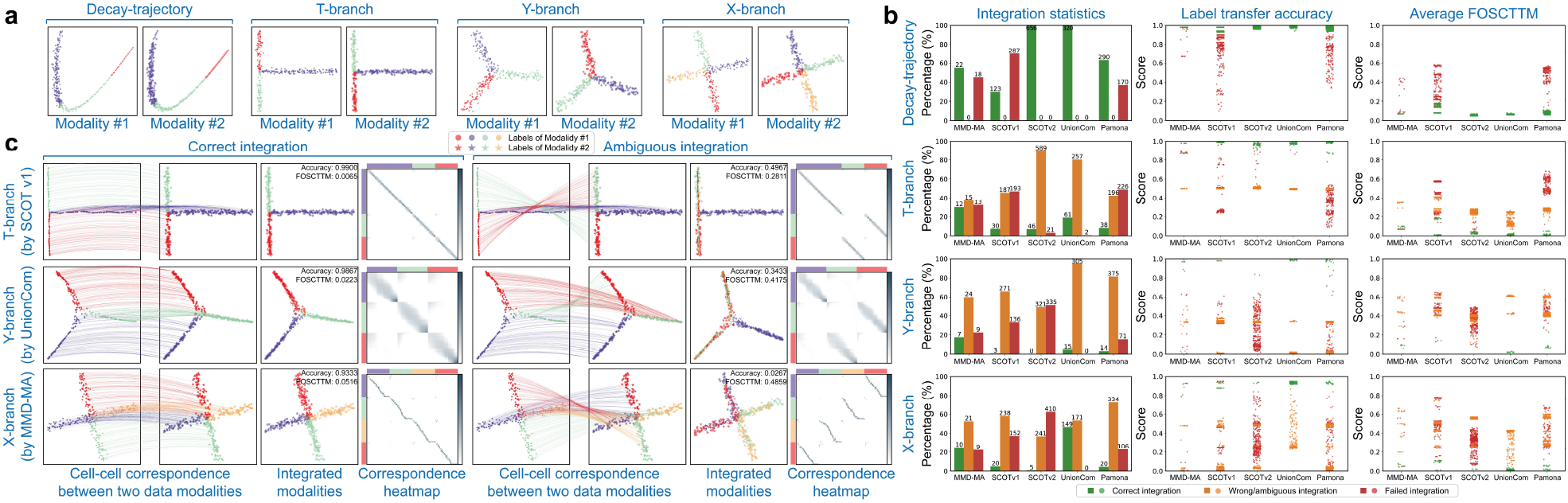
Ambiguous integrations are revealed on simulated datasets. (**a**) Four simulated datasets were constructed, with one dataset (*i*.*e*., Decay-path) having no ambiguity, while the other three exhibit branch-level ambiguity. The cell labels are indicated by color, where branches of the same color between two modalities should be aligned. Different modalities are represented by different markers. (**b**) Ambiguous mappings occur universally across mainstream diagonal integration methods, quantified using the LTA and average FOSCTTM metrics. (**c**) Ambiguous mappings can be qualitatively detected by examining the cross-modality cell-cell correspondence matrix generated by existing integration methods, which forms the basis of SONATA’s intuition.

Alongside the simulated datasets, We employed three real multimodal single-cell datasets, each with co-assayed cell-cell correspondence information, to uncover the artificial integration issues. The sc-GEM dataset (Fig. 3**a**) was generated using the sc-GEM technology [30], a sequencing technology that simultaneously profiles gene expression and DNA methylation states within the same cell. The dataset was derived from human somatic cell samples undergoing conversion to induced pluripotent stem (iPS) cells [30], which contains *n*_*x*_ = *n*_*y*_ = 177 cells with *p*_*x*_ = 34 dimensions of gene expression features and *p*_*y*_ = 27 dimensions of chromatin accessibility features, respectively.

**Figure 3.**
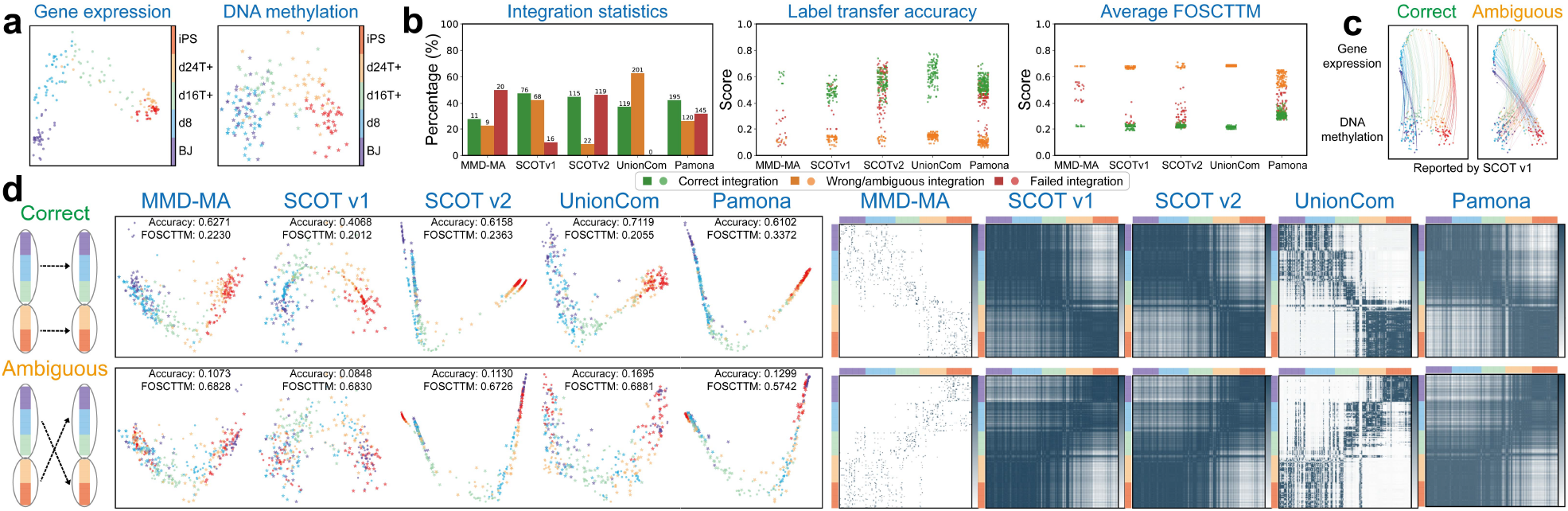
Ambiguous integrations are revealed on sc-GEM dataset. (**a**) The sc-GEM dataset consists of two data modalities: gene expression and DNA methylation. (**b**) Ambiguous mappings occur universally across mainstream diagonal integration methods. (**c**) Ambiguous mappings occur when one modality is reversely aligned to the other during the integration process. (**d**) Ambiguous mappings can be qualitatively revealed through the aligned manifold and the crossmodality cell-cell correspondences.

The SNARE-Seq dataset (Fig. 4**a**) was generated using the SNARE-seq technology [31], a sequencing technology that simultaneously profiles gene expression and chromatin accessibility within the same cell. The dataset was derived from a mixture of BJ, H1, K562, and GM12878 human cell lines [31], which contains *n*_*x*_ = *n*_*y*_ = 1047 cells in both modalities. Following [13], we carried out dimensionality reduction on the chromatin accessibility information using the topic modeling framework cisTopic [32], resulting in *p*_*x*_ = 19 dimensions of chromatin accessibility features. We also performed dimensionality reduction on the gene expression information using PCA, resulting in *p*_*y*_ = 10 dimensions of gene expression features.

**Figure 4.**
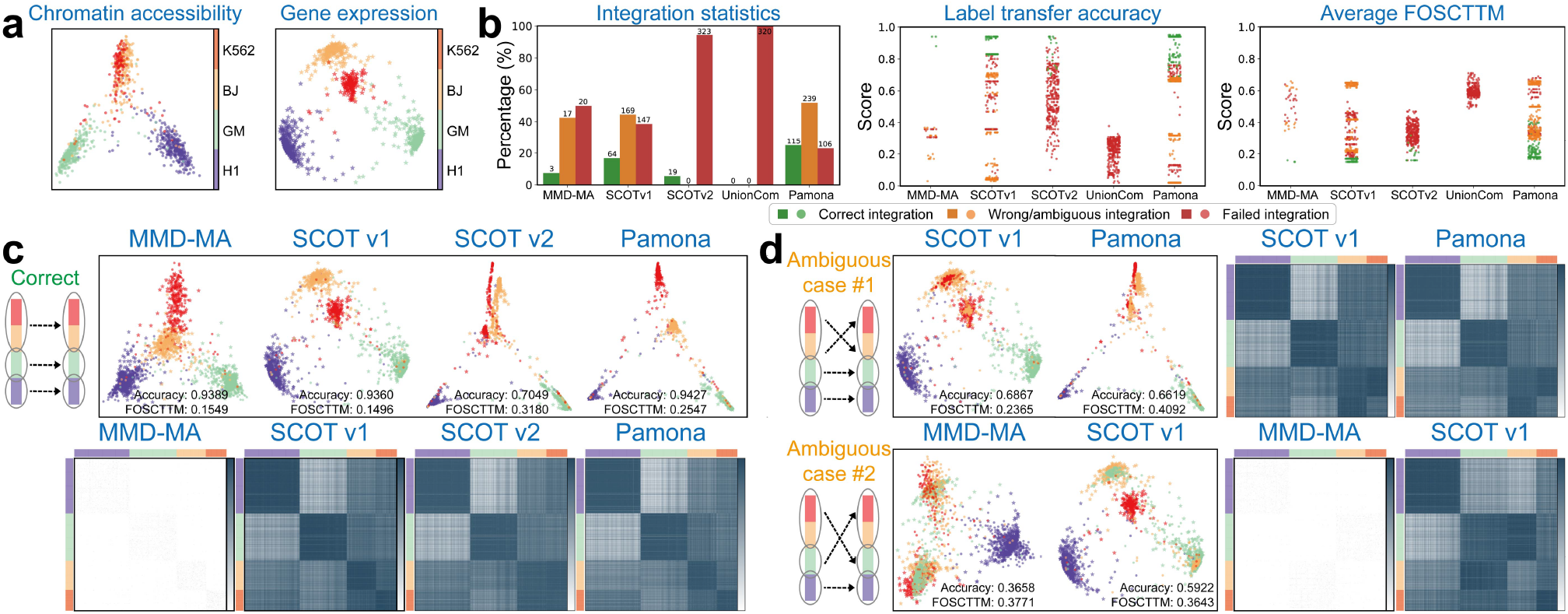
Ambiguous integrations are revealed on SNARE-Seq dataset. (**a**) The dataset consists of two data modalities: chromatin accessibility and gene expression. (**b**) Ambiguous mappings occur across most diagonal integration methods, except for UnionCom, which always fails regardless of parameter settings. (**c**) Correct integration accurately aligns cells of the same type across modalities. (**d**) Artificial integrations can result from different types of ambiguous mappings.

The sc-NMT (Fig. 5**a**) was generated using the sc-NMT sequencing technology [33], a sequencing technology that simultaneously profiles DNA methylation and chromatin accessibility within the same cell. The dataset was collected from mouse gastrulation at three-time: embryonic day 5.5 (E5.5), E6.5 and E7.5. Following [15], we filtered out cells where all features were marked as missing values, resulting in *n*_*x*_ = 612 cells in chromatin accessibility and *n*_*y*_ = 709 cells in DNA methylation. A UMAP-based dimensionality reduction [34] was then performed on each dataset separately, yielding a dimensionality of *p*_*x*_ = *p*_*y*_ = 300 for both.

**Figure 5.**
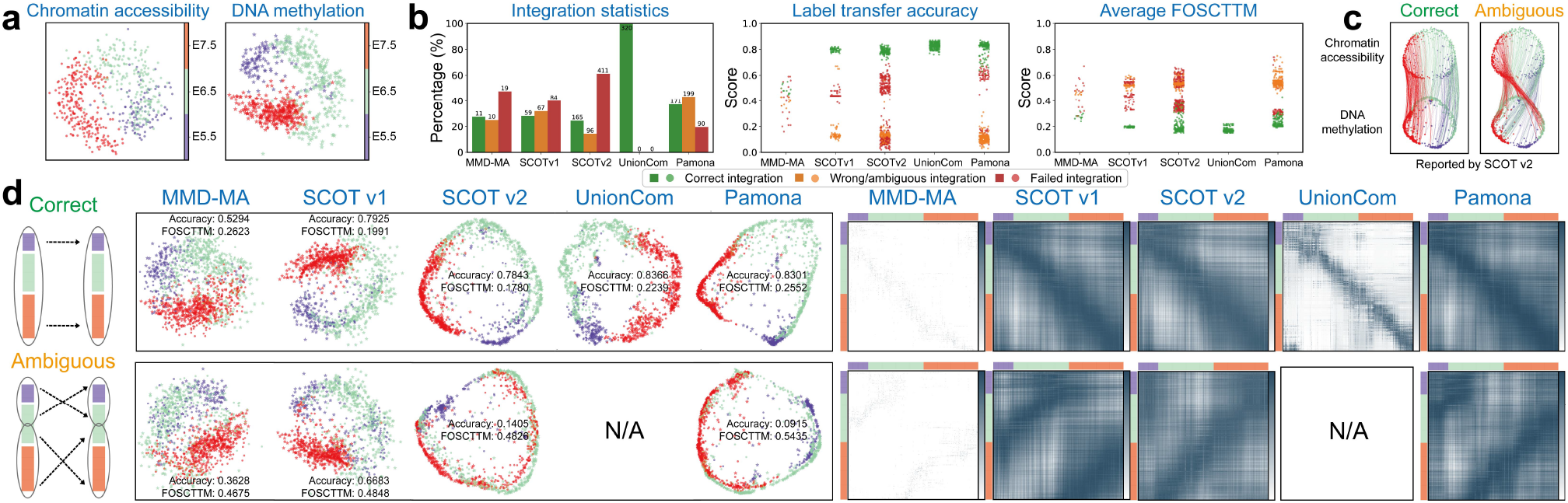
Ambiguous integrations are revealed on sc-NMT dataset. (**a**) The dataset consists of two data modalities: chromatin accessibility and DNA methylation. (**b**) Ambiguous mappings occur across most diagonal integration methods, except for UnionCom, which always succeeds across different settings. (**c**) Ambiguous mappings occur when two parts of one modality are reversely aligned to their counterpart, respectively. (**d**) Ambiguous mappings can be qualitatively revealed through the aligned manifold and the cross-modality cell-cell correspondences.

### 3.2 Experimental settings

We investigated the occurrence of artificial integrations caused by ambiguous mapping across various diagonal integration methods. To demonstrate the universality of this issue, we benchmarked several mainstream diagonal integration methods: MMD-MA [11], SCOT [13, 14], UnionCom [15], and Pamona [16]. We investigated two widely used versions of SCOT: SCOTv1 [13] and SCOTv2 [14], with the latter extending the former by explicitly addressing disproportionate cell types.

We utilized two metrics to quantitatively assess the performance of multimodal single-cell diagonal data integration. Since all simulated and real datasets include cell-cell correspondence information, we employed the average “Fraction Of Samples Closer Than the True Match” (FOSCTTM) metric [11] to quantify how well the integration preserves cell-cell correspondence. Specifically, for each cell in one modality, FOSCTTM calculates the fraction of cells in the other modality that are closer to it than its true match. We then compute the average FOSCTTM by taking the mean of the FOSCTTM values across all cells in both modalities. Since the average FOSCTTM measures the proportion of falsely matched cells, it ranges from 0 to 1, with smaller values indicating better integration performance. Additionally, since both simulated and real datasets contain cell label information, we utilized “Label Transfer Accuracy” (LTA) [15] to assess how well the shared cell labels can be transferred from one modality to another following the integration. Specifically, LTA evaluates the accuracy of transferring cell labels from one modality to another, based on their neighborhood relationships in the aligned modality. Since LTA measures the proportion of correctly predicted labels, it ranges from 0 to 1, with larger values indicating better integration performance.

## 4 Results

### 4.1 Ambiguous integrations are revealed through simulated data analysis

We first highlighted the artificial integrations arising from ambiguous mapping in simulated datasets (Fig. 2**a**). We employed five mainstream diagonal integration methods to integrate the two modalities of each dataset, utilizing a diverse range of parameter settings recommended by each method. (Refer to A.1 for the detailed parameter settings). Surprisingly, ambiguous mappings occur universally across all these integration methods and are more prevalent than correct mappings (Fig. 2**b**). In other words, blindly trusting the integration results may easily mislead users into pursuing hypotheses based on inaccurate findings.

We then examined the ambiguous mappings both qualitatively and quantitatively. For each dataset, we utilized Principal Component Analysis (PCA) to project the two aligned modalities into a shared 2D space for visualization (Fig. 2**c**). In the three datasets exhibiting ambiguities, all diagonal integration methods align the branches without grouping those with the same label together. For example, in the T-shaped branch dataset, the green branch in the first modality is incorrectly mapped to the red branch in the second modality, and vice versa. Note that ambiguous mappings can be detected through the cross-modality cell-cell correspondence matrix generated by existing integration methods. In correct integrations, this matrix exhibits a block-diagonal structure, while artificial integrations disrupt this structure by permuting the blocks. Detailed ambiguous results from all five methods are provided in A.5.

### 4.2 Artificial integrations are identified through real data analysis

We then highlighted the artificial integrations arising from ambiguous mapping in real datasets. We began by examining the sc-GEM dataset, which comprises two data modalities: chromatin accessibility and DNA methylation (Fig. 3**a**). Both modalities display similar trajectories with the same order of cell types, transitioning from BJ (human foreskin fibroblast) cells at one end to iPS cells at the other, reflecting the underlying biological process. We employed five diagonal integration methods to integrate the two modalities of each dataset with a diverse range of parameter settings (Fig. 3**b**). Surprisingly, ambiguous mappings occur universally across all these integration methods when one modality is reversely aligned to the other during the integration process. Specifically, based on the cell type annotations, iPS cells, located initially at only one end of the trajectory, now reside in both ends of the aligned trajectories (Fig. 3**c**). The reverse alignment can be revealed qualitatively by the aligned manifold and the cross-modality cell-cell correspondence matrix generated by existing integration methods (Fig. 3**d**).

We next examined the SNARE-Seq dataset, which comprises two data modalities: gene expression and chromatin accessibility (Fig. 4**a**). Both modalities are derived from a mixture of BJ, H1, K562, and GM12878 human cell lines, with H1, BJ, and K562 cells clustered in different groups. We employed five diagonal integration methods to integrate the two modalities of each dataset with a diverse range of parameter settings (Fig. 4**b**). Surprisingly, ambiguous mappings occur across most diagonal integration methods, except for UnionCom, which always fails regardless of parameter settings. It is important to note that ambiguous mappings may not be unique (Fig. 4**d**). For example, one type of ambiguous mappings confuses the correspondence between the BJ cluster and the K562 cluster, while the other type flips the correspondence between the GM cluster and the aggregated BJ/K562 clusters.

Lastly, we examined the sc-NMT dataset, which comprises two data modalities: DNA methylation and chromatin accessibility (Fig. 5**a**). Both modalities exhibit a similar cyclical trajectory, reflecting the same temporal order of cell collection, transitioning from E5.5 to E6.5 and then to E7.5. We employed five diagonal integration methods to integrate the two modalities of each dataset with a diverse range of parameter settings (Fig. 5**b**). Surprisingly, ambiguous mappings occur in most diagonal integration methods, with the exception of UnionCom, which consistently succeeds across various settings. The ambiguous mapping occurs when two segments of one modality (E5.5 and part of E6.5, and E7.5 and part of E6.5) are reversely aligned to their corresponding counterparts (Fig. 5**c**). The ambiguous mapping can be revealed qualitatively by the cross-modality cell-cell correspondences generated by existing integration methods (Fig. 5**d**).

### 4.3 SONATA consistently diagnoses and improves artificial integrations

Given the struggling performance of existing integration methods in both real and simulated datasets, we investigated whether SONATA could assist these methods in identifying ambiguities within the datasets, thereby improving the integration results. As shown in Fig. 6**a**, SONATA accurately identifies ambiguous cell groups between branches that exhibit geometric resemblance in the simulated datasets. For example, SONATA identifies geometrically similar cell pairs between two and three branches with resemblance in the T-shaped and Y-shaped branch datasets, respectively. The identified ambiguous cell groups are consistent across different modalities, demonstrating SONATA’s robustness in detecting ambiguity regardless of the modality.

**Figure 6.**
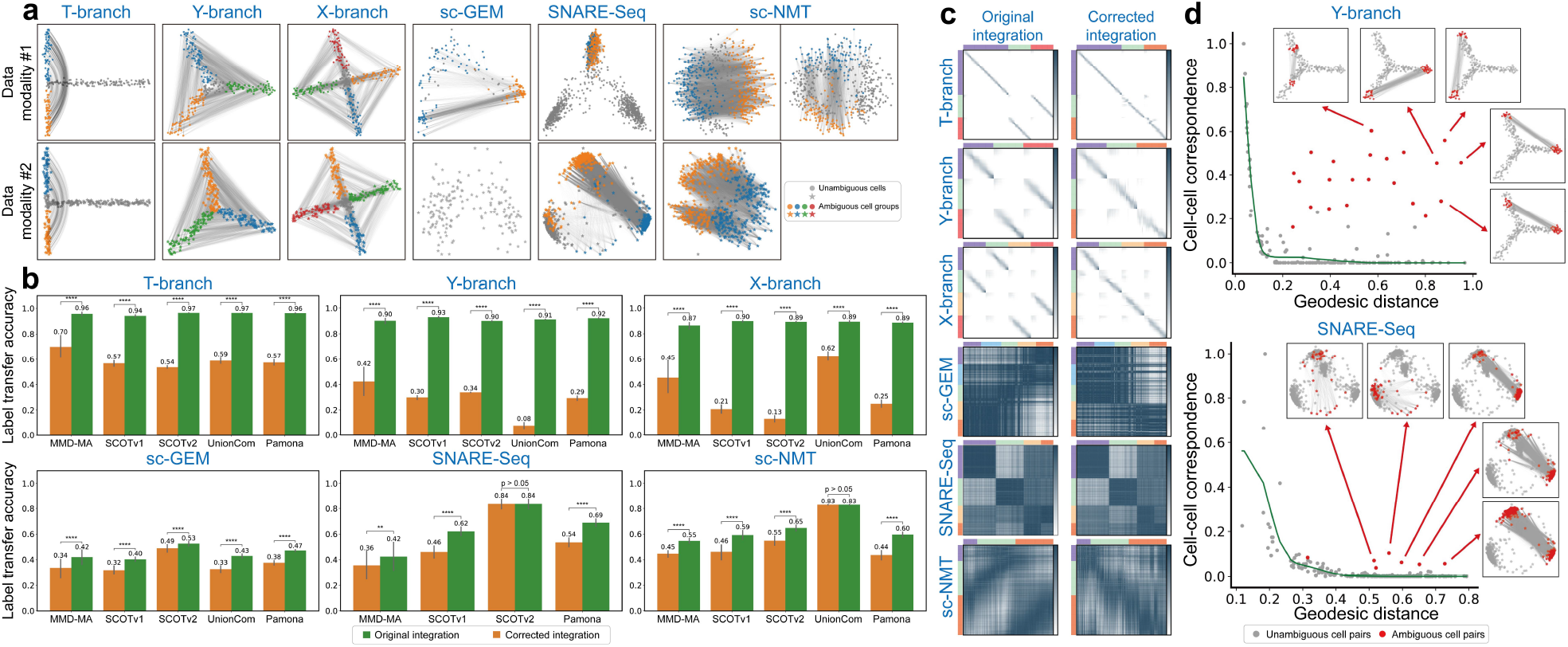
SONATA secures diagonal integrations in both simulated and real datasets. (**a**) SONATA identifies ambiguous cell groups that may cause confusion during the integration process. (**b**) SONATA generates alternative integration solutions that are often overlooked by existing methods, among which the best solution demonstrates significantly improved integration performance. For scientific rigor, the performance comparison between the original and the best alternative solution is quantified using one-tailed two-sample t-tests to calculate p-values: * * * * p-value ≤ 0.0001; * * *p-value ≤ 0.001; * * p-value ≤ 0.01; *p-value ≤ 0.05. (**c**) The improved integration performance can be qualitatively observed through the changes in cross-modality cell-cell correspondences. (**d**) SONATA identifies ambiguous cell-cell mappings using a statistical confidence measure derived from their observed correspondences.

In three real datasets, SONATA demonstrates distinct behaviors across different datasets. In the sc-GEM dataset, SONATA identifies the ambiguity between cells at one end of the trajectory (iPS and d24T+ cells) and cells at the opposite end (BJ and d8 cells). The identified ambiguous cell groups are consistent with the observed reverse cross-modality alignment found using five diagonal integration methods (Fig. 3). In the SNARE-Seq dataset, SONATA identifies different ambiguous cell groups across different data modalities. Specifically, in the chromatin accessibility modality, SONATA identifies ambiguous cell groups between the BJ and K562 clusters, while in the gene expression modality, it detects ambiguous groups between the GM cluster and the BJ/K562 clusters. It is important to note that we identified two distinct types of ambiguous mappings while applying five diagonal integration methods for data integration. These mappings correspond precisely to the two ambiguous cell groups reported by SONATA. In the sc-NMT dataset, SONATA identifies different ambiguous cell groups even within a single data modality. Specifically, in the chromatin accessibility modality, SONATA identifies two types of ambiguous cell groups. One type of ambiguous cell group maps E5.5 and a portion of E6.5 to E7.5, while the other type maps E6.5 to sections of both E5.5 and E7.5. Given that the modality follows a cyclical trajectory, E5.5, E6.5, and E7.5 display geometric similarities to one another, making both types of ambiguous cell groups justifiable.

The presence of ambiguous cell groups suggests the existence of alternative yet overlooked integration solutions. Therefore, SONATA leverages these ambiguous cell groups to generate alternative integration solutions that are often overlooked by existing methods. To determine whether SONATA truly uncovers the previously overlooked integrations, we examined whether the alternative solutions lead to improved integration performance quantitatively. As shown in Fig. 6**b**, the best alternative solution identified by SONATA consistently and significantly outperforms the solutions provided by all existing integration methods. The improved integration performance can be qualitatively observed through the changes in cross-modality cell-cell correspondences (Fig. 6**c**). It is important to note that the detection of ambiguous cell-cell mappings is interpretable, as statistically confident mappings can be qualitatively visualized for users (Fig. 6).

## 5 Discussion and conclusion

In this study, we propose SONATA, a novel diagnostic method for diagonal data integration of multimodal single-cell data that aims to identify potential artificial integrations resulting from ambiguous mapping. The key novelty of SONATA is twofold. First and foremost, we demonstrate that artificial integrations resulting from ambiguous mapping in diagonal data integration are widespread yet surprisingly overlooked, occurring across all mainstream diagonal integration methods. Additionally, we define a novel cell-cell ambiguity measurement, assessed through a statistical confidence measure for each pair of cells within the same modality. This ambiguity is then used to identify cell groups with similar geometric contexts, which could potentially lead to artificial integrations in cross-modality mapping.

Unlike existing methods that recklessly report an arbitrary integration solution, SONATA distinguishes whether the integration is unique or if multiple alternative solutions exist, prompting users to carefully discern the true biological solution before proceeding to downstream analysis. It is worth noting that SONATA is not designed to replace any existing pipelines for diagonal data integration; instead, SONATA works simply as an add-on to an existing pipeline for achieving more reliable integration. We applied SONATA to both simulated and real multimodal single-cell datasets to demonstrate its empirical utility. Our experiments on real datasets highlight SONATA’s ability to safeguard diagonal integration of gene expression, DNA methylation, and chromatin accessibility modalities against ambiguous mapping from mainstream methods, effectively informing potential users where these methods might fail.

This study points to several promising directions for future research. SONATA is designed as a post-processing method designed to complement existing manifold alignment methods. We rationalize that the proposed cell-cell ambiguity measurement can be re-purposed to measure the cell-cell similarity between different modalities. Using SONATA directly for both manifold alignment and disambiguation would be interesting directions to pursue. Additionally, several critical single-cell applications, such as batch-effect correction [35] and comparative analysis between case-control studies [36], are closely associated with the manifold alignment. We would like to use SONATA to identify and resolve the ambiguity in wide applications.

## A Appendix

### A.1 Baseline settings

To evaluate baseline performance on different datasets, we tested the parameters within the recommended ranges as described in the original publications. For SCOTv1 and SCOTv2, we followed the SCOT tutorial, testing the parameter *k* between [20, *n/*5], where *n* denotes the number of samples (cells) in the smallest dataset. Additionally, we followed the tutorial’s guidance when testing the range of the coefficient of the entropic regularization term *e*. For UnionCom, we tested *k* and *ρ* based on the robustness analysis outlined in the original work. In the case of MMD-MA, we preprocessed the input data using the linear kernel as described in the literature, and followed the method in [37] to automatically compute the bandwidth parameter *σ*. We experimented with two sets of *λ*_1_ and *λ*_2_ for each dataset, conducting tests across 20 random seeds for each setting.

The detailed parameter settings are provided in Table A.1, A.2.

**Table A.1:** Hyperparameter settings for baselines.

| Method | Fixed Parameters | Tested Parameters |
| --- | --- | --- |
| MMD-MA | $num\_feat = 4$,<br>$epoch = 5000$ | $\lambda_1, \lambda_2 \in [(1e-3, 1e-3), (1e-6, 1e-7)]$ for Decay-trajectory<br>$\lambda_1, \lambda_2 \in [(1e-3, 1e-3), (1e-6, 1e-6)]$ for other simulated datasets<br>$\lambda_1, \lambda_2 \in [(1e-3, 1e-3), (1e-6, 1e-6)]$ for sc-GEM<br>$\lambda_1, \lambda_2 \in [(1e-3, 1e-3), (1e-5, 1e-5)]$ for SNARE-Seq<br>$\lambda_1, \lambda_2 \in [(1e-3, 1e-3), (1e-5, 1e-6)]$ for sc-NMT<br>$random\_seed \in [0, 1, 2, \dots, 19]$ for all datasets |
| SCOT v1 | Default Settings | $k \in [20, 21, 22, \dots, 60]$ for simulated datasets<br>$k \in [20, 21, 22, \dots, 35]$ for sc-GEM<br>$k \in [20, 25, 30, \dots, 205]$ for SNARE-Seq<br>$k \in [20, 25, 30, \dots, 120]$ for sc-NMT<br>$e \in [1e-3, 5e-3, 1e-2, 5e-2, 1e-1]$ for all datasets<br>Tested Settings:<br>1. **mode=distance, metric=euclidean**, for all datasets<br>2. **mode=connectivity, metric=correlation** for all datasets |
| SCOT v2 | Default Settings | $k \in [20, 21, 22, \dots, 60]$ for simulated datasets<br>$k \in [20, 21, 22, \dots, 35]$ for sc-GEM<br>$k \in [20, 25, 30, \dots, 205]$ for SNARE-Seq<br>$k \in [20, 25, 30, \dots, 120]$ for sc-NMT<br>$e \in [1e-2, 5e-2, 1e-1]$ for SNARE-Seq<br>$e \in [5e-3, 1e-2, 5e-2, 1e-1]$ for simulated, sc-GEM and SNARE-Seq<br>$\rho \in [1e-3, 1e-2, 1e-1, 1.0]$ for all datasets<br>Tested Settings:<br>1. **mode=distance, metric=euclidean** for all datasets,<br>2. **mode=connectivity, metric=correlation** for sc-NMT |

**Table A.2:** Hyperparameter settings for baselines.

| Method | Fixed Parameters | Tested Parameters |
| --- | --- | --- |
| UnionCom | Default Settings | $k \in [5, 6, 7, \dots, 20]$ for all datasets<br>$\rho \in [1, 2, 3, \dots, 20]$ for all datasets |
| Pamona | Default Settings | $k \in [5, 6, 7, \dots, 50]$ for all datasets<br>$e \in [1e - 3, 5e - 3, 1e - 2, 5e - 2, 1e - 1]$ for all datasets<br>Tested Settings:<br>1. `mode=distance`, `metric=minkowski` for all datasets,<br>2. `mode=connectivity`, `metric=correlation` for all datasets |

### A.2 Details of SONATA

#### A.2.1 Acceleration of spline fitting

SONATA fits a cubic smoothing spline to model the probability of cell-cell correspondence as a function of geodesic distance. However, fitting a spline directly to all cell-cell pairs may not be ideal due to the computational expense of handling the quadratic number of pairs, and the noise they introduce could negatively impact the fitting performance. To achieve a smooth and efficient spline fit, we follow the intuition that ambiguous cells are unlikely to exist in isolation but are more likely to appear alongside their neighboring cells within the data manifold. Thus, we partition the cells into approximately 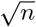 groups using hierarchical clustering, where *n* is the total number of cells in the dataset (Refer to Sec.A.3.2 for the guidance on selecting the number of groups.). We then use the average correspondence and geodesic distance between the cells in each pair of groups to perform the fit.

#### A.2.2 Generating alternative integration solutions

SONATA detects ambiguous cell groups which suggest the existence of alternative yet overlooked integration solutions. The rationale for these ambiguous cell groups is that one group can be substituted for another based on their geometric resemblance during cross-modality integration. Specifically, for two ambiguous group of cells, denoted as *G*_*s*_, *G*_*t*_ ⊆ {1, 2, , *n*}, we obtain their cell mapping 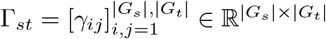 from the self-alignment conducted in Sec. 2.2. Thereby, given an existing diagonal integration solution in the form of cross-modality correspondence matrix 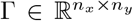, SONATA reports an alternative solution 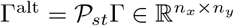 where the permutation matrix 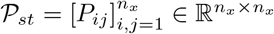 is defined as:

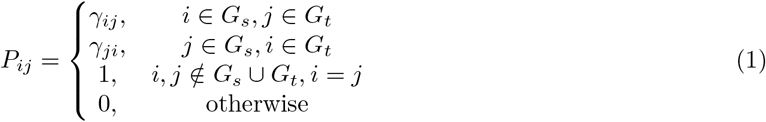

### A.3 Parameter Settings in SONATA

#### A.3.1 Noise Level

SONATA introduces random noise to generate variational versions of itself. This process enables alignment with a slightly perturbed manifold, which helps prevent trivial mappings. Instead of aligning cells with their exact copies, it encourages them to align with geometrically similar cells across subtly transformed versions of the data. Conceptually, the noise level should neither be too low, which could result in trivial solutions, nor too high, which could disrupt the manifold’s integrity. We propose that, regardless of the noise type applied, a noise level must be established that preserves the manifold’s structural integrity. Consequently, the optimal noise level may vary depending on the specific dataset. To quantify this, we calculated the Pearson correlation between the geodesic matrices of the original dataset 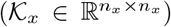 and the noisy dataset 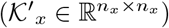 at various levels of Gaussian noise.

As shown in Fig. A.1, different datasets exhibit varying sensitivity to noise levels. For example, in real datasets with clear structures, such as SNARE-Seq, which contains several well-separated cell groups, higher levels of noise can be tolerated without significant disruption to the underlying structure. Conversely, in datasets like sc-NMT, which display a circular-like structure, the introduction of noise is more likely to alter this structure adversely. We recommend maintaining a minimum of 80% structural similarity, regardless of the noise type employed. This threshold was established in our experiments to ensure the robustness of the manifold.

**Figure A.1:**
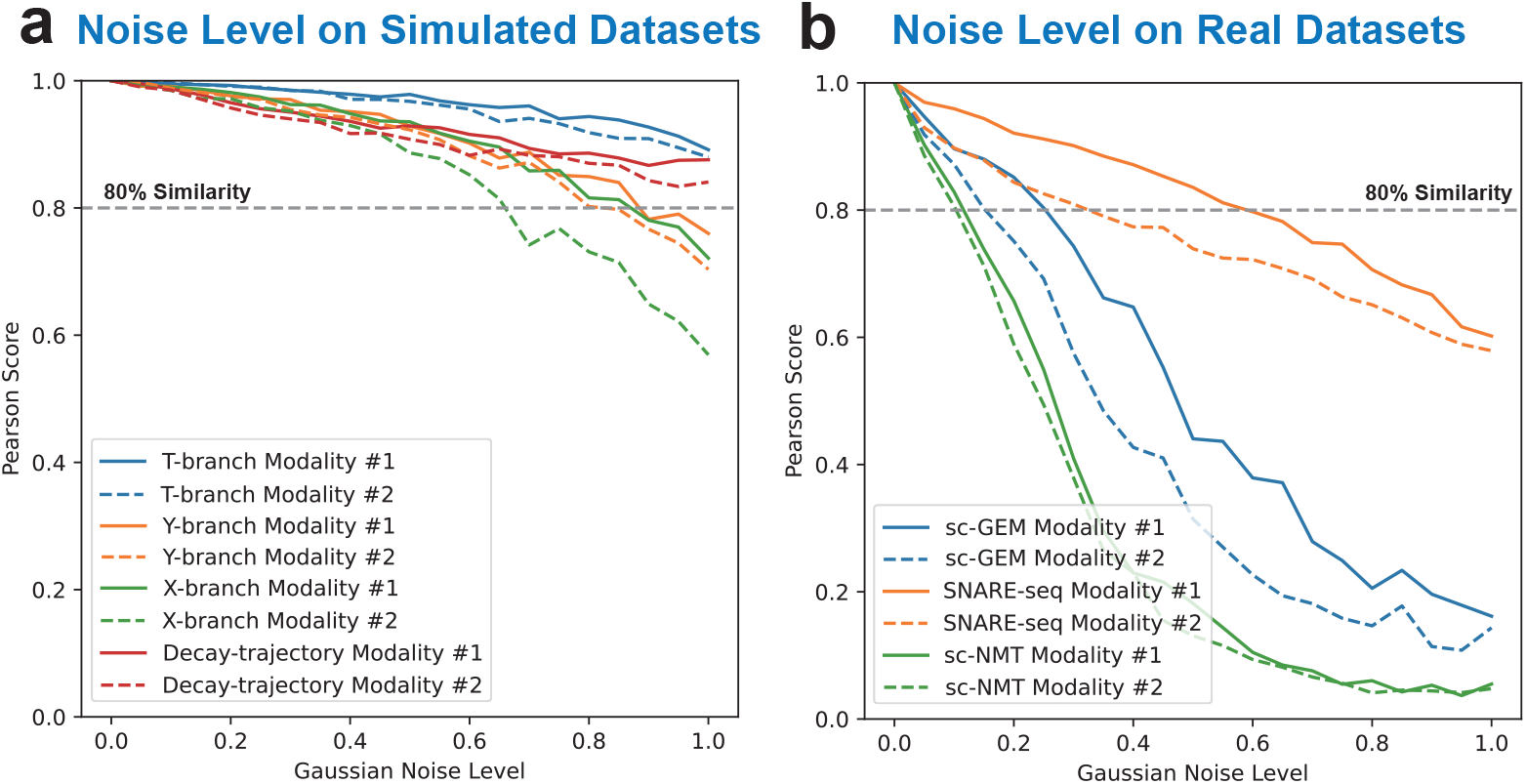
Gaussian Noise Level.

#### A.3.2 Number of Groups in Clustering

We evaluated the performance of SONATA with respect to the number of clusters *k* in the clustering process, as shown in Fig. A.2. We used ambiguous cases from the sc-GEM dataset, integrated by SCOTv1 (**Fig. 3c**), as the test case and varied *k* to observe SONATA’s impact on Label Transfer Accuracy.

From Fig. A.2**a**, we observe that within the recommended range of, 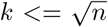 where *n* = 177 represents the sample size), SONATA demonstrates consistently high performance in Label Transfer Accuracy, maintaining stability across the tested values of *k*.

However, when *k* exceeds this threshold, performance begins to degrade. This decline is attributed to the fact that a disproportionately large number of clusters can lead to imbalanced clustering, introducing noise into the results. For instance, in Fig. A.2**b**, when *k*=7, the clustering is reasonable, and the ambiguous groups are effectively identified. In contrast, when *k* increases to 15, clusters 9 and 14 contain only 2 and 3 samples, respectively, which undermines the reliability of these clusters to represent distinct groups (Fig. A.2**c**).

Therefore, we recommend setting *k* to values below 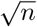 to avoid creating clusters with very few samples, ensuring the robustness of the results.

**Figure A.2:**
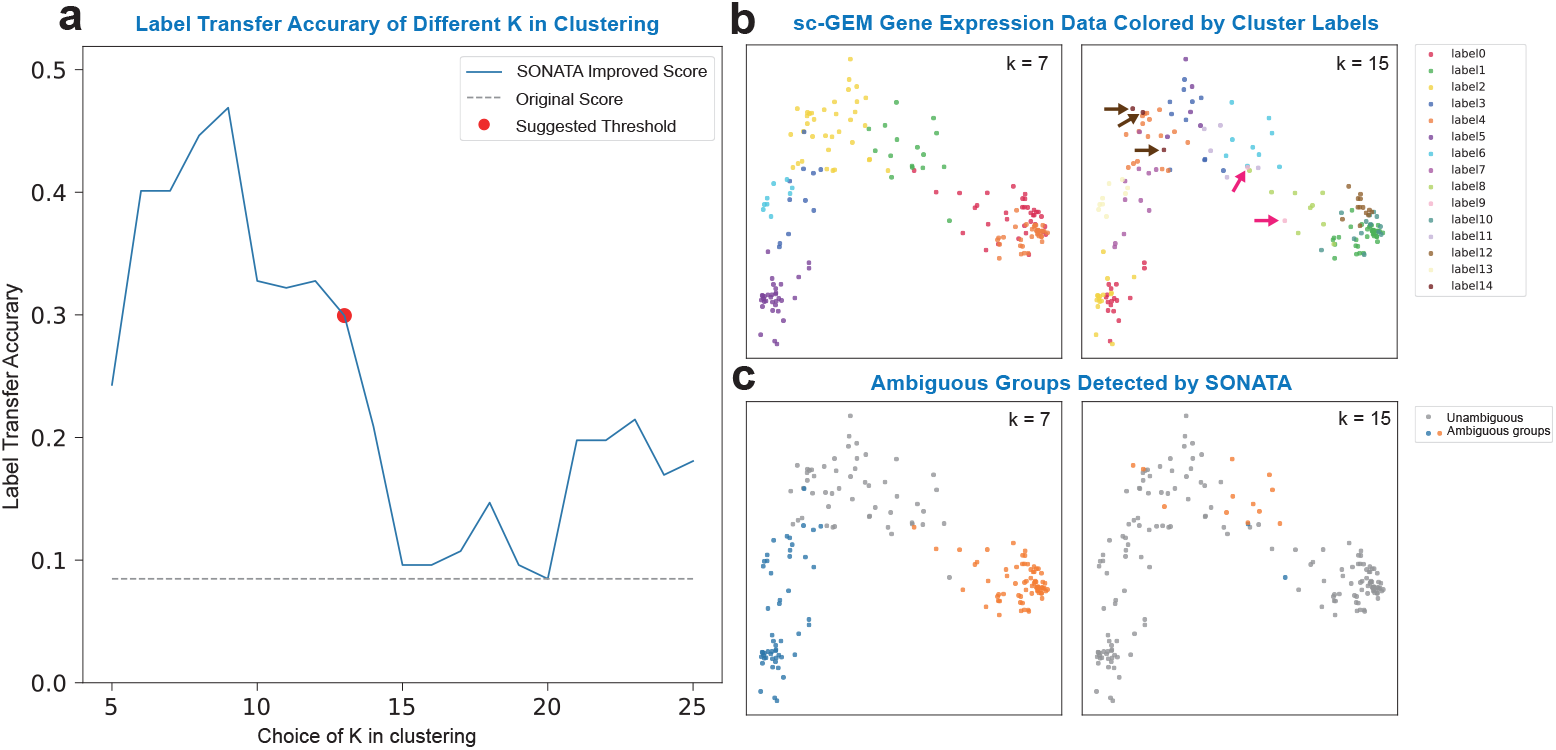
Cluster Number Settings in SONATA.

### A.4 Simulated data analysis

**Figure A.3:**
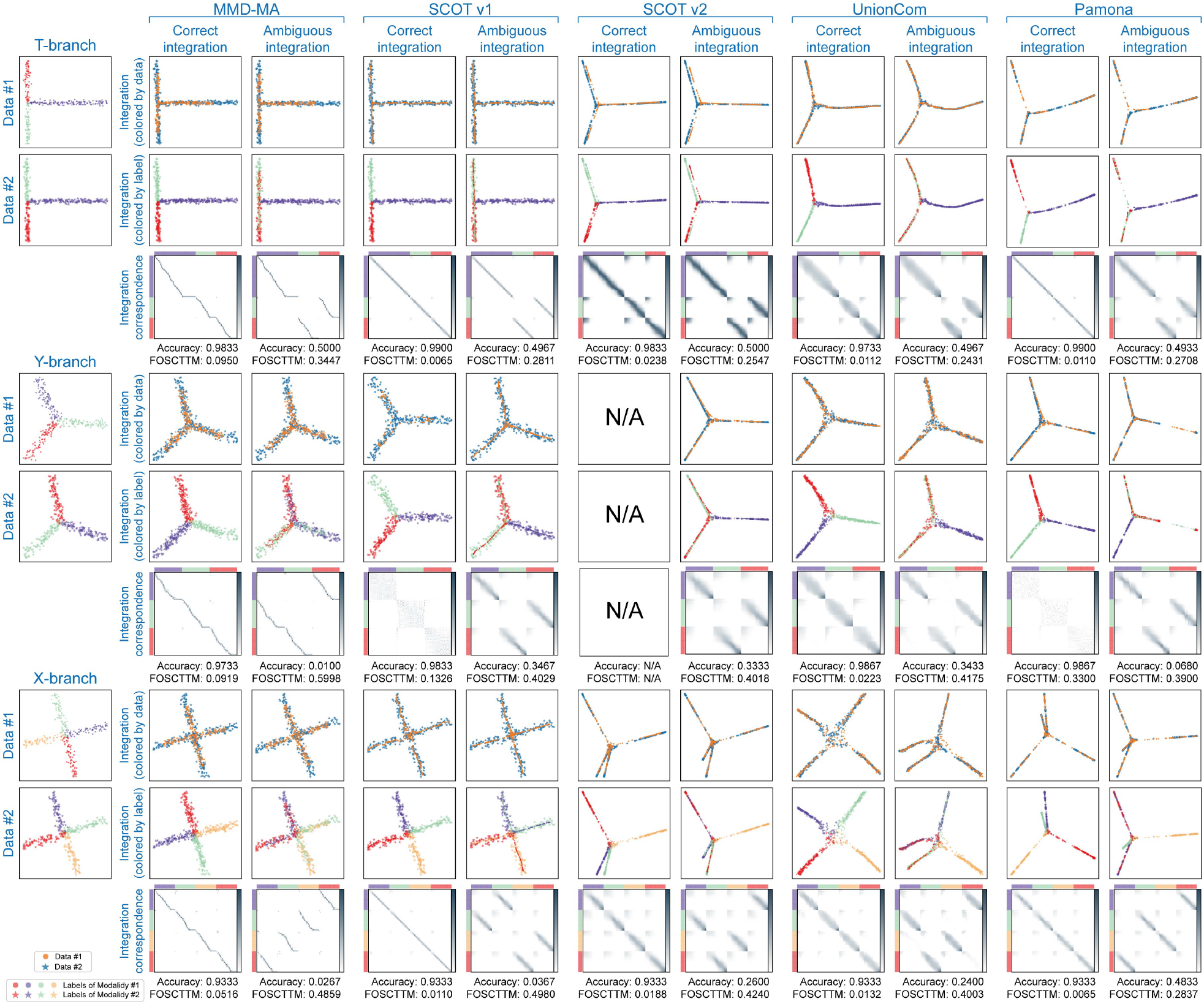
Comprehensive ambiguous integration results on simulated datasets.

**Figure A.4:**
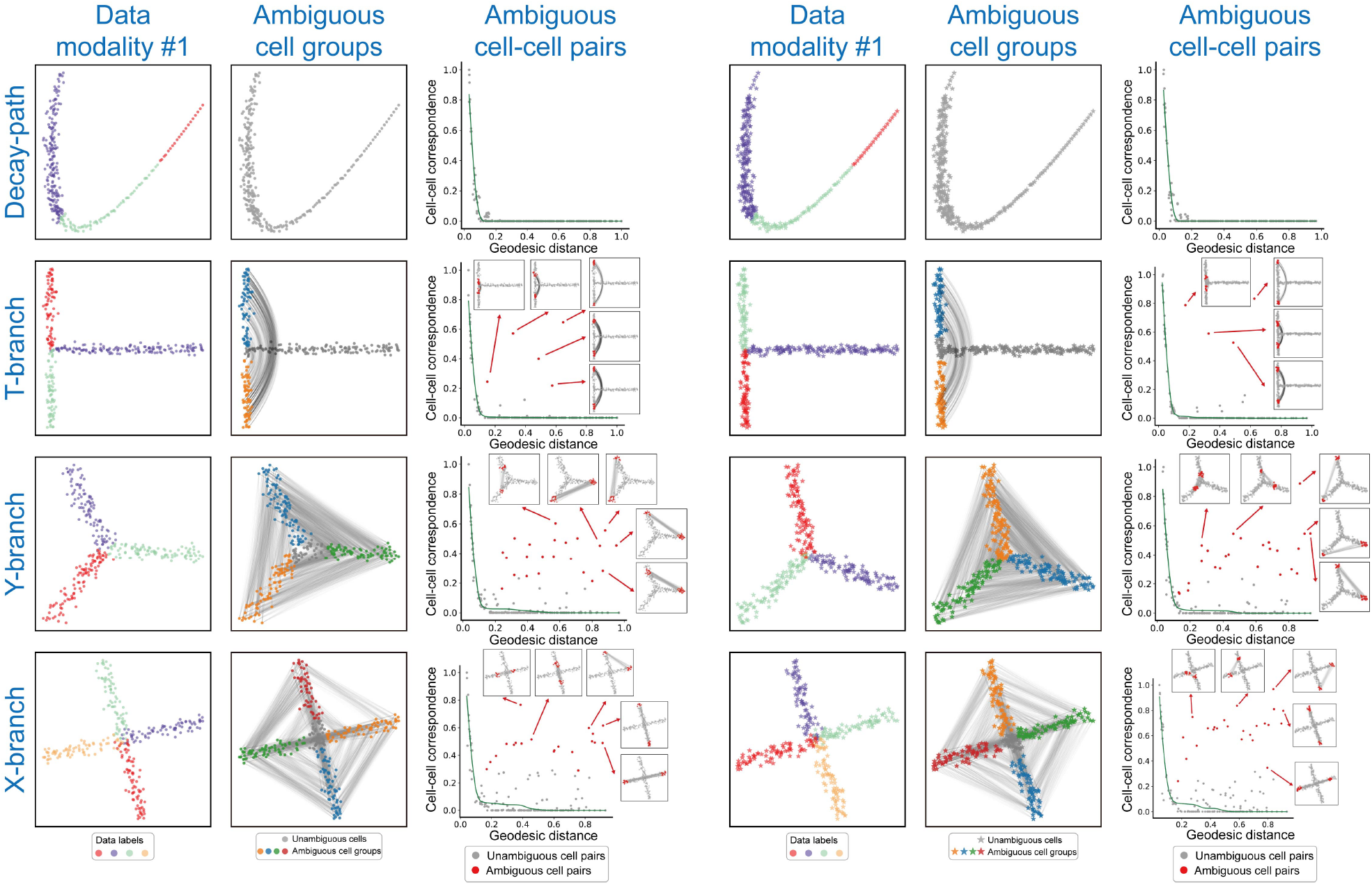
Comprehensive SONATA results on simulated datasets.

#### A.5 Real data analysis

**Figure A.5:**
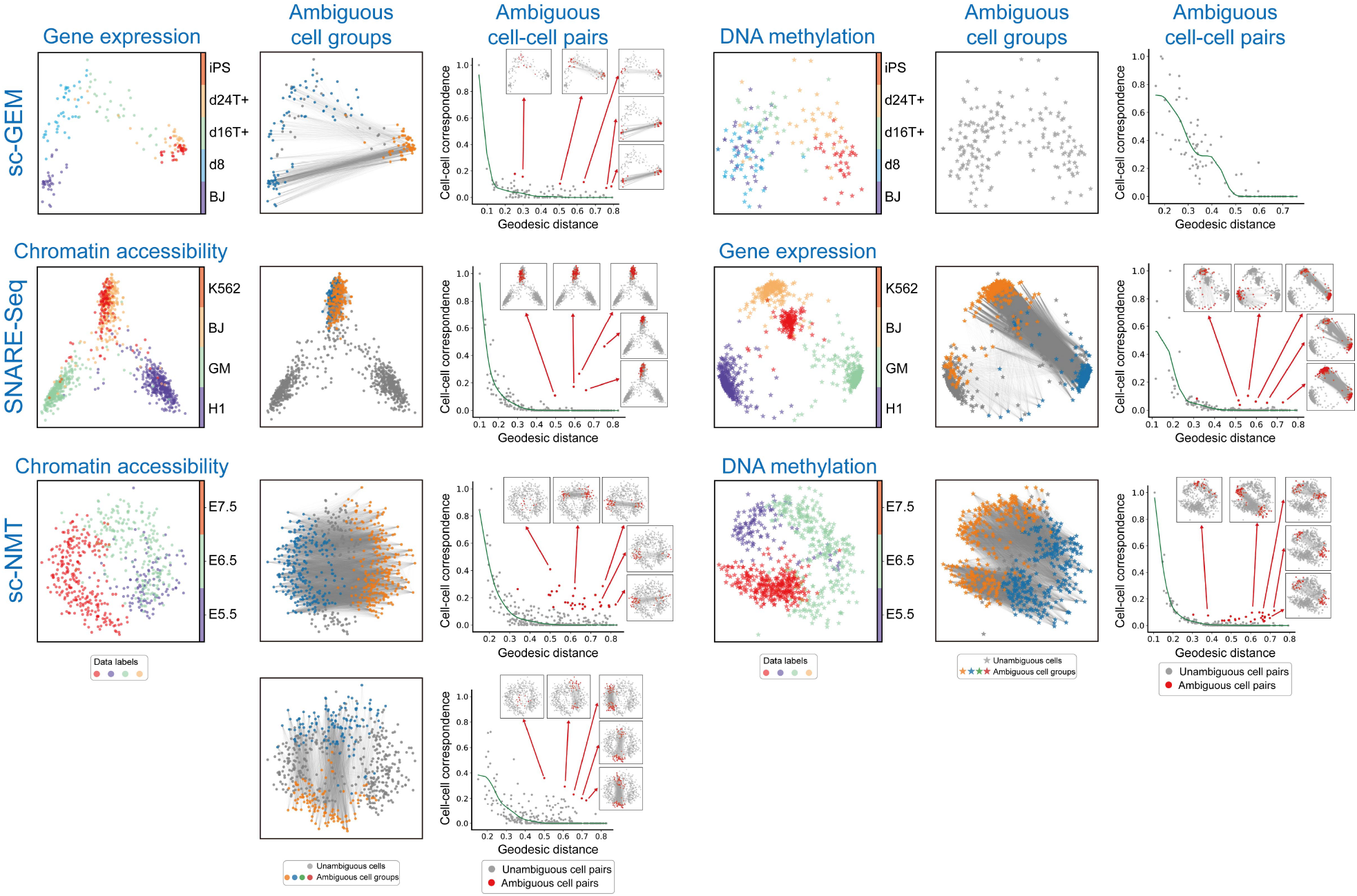
Comprehensive SONATA results on real datasets.

## Footnotes

1 The Apache licensed source code of SONATA is available at https://github.com/batmen-lab/SONATA.

